## Supplemental Figures for "A multiplexed, single-cell sequencing screen identifies compounds that increase neurogenic reprogramming of murine Muller glia"

**This PDF file includes:**

Figures S1 to S3
Table S1

**Other supporting materials for this manuscript include the following:**

Datasets S1 to S3

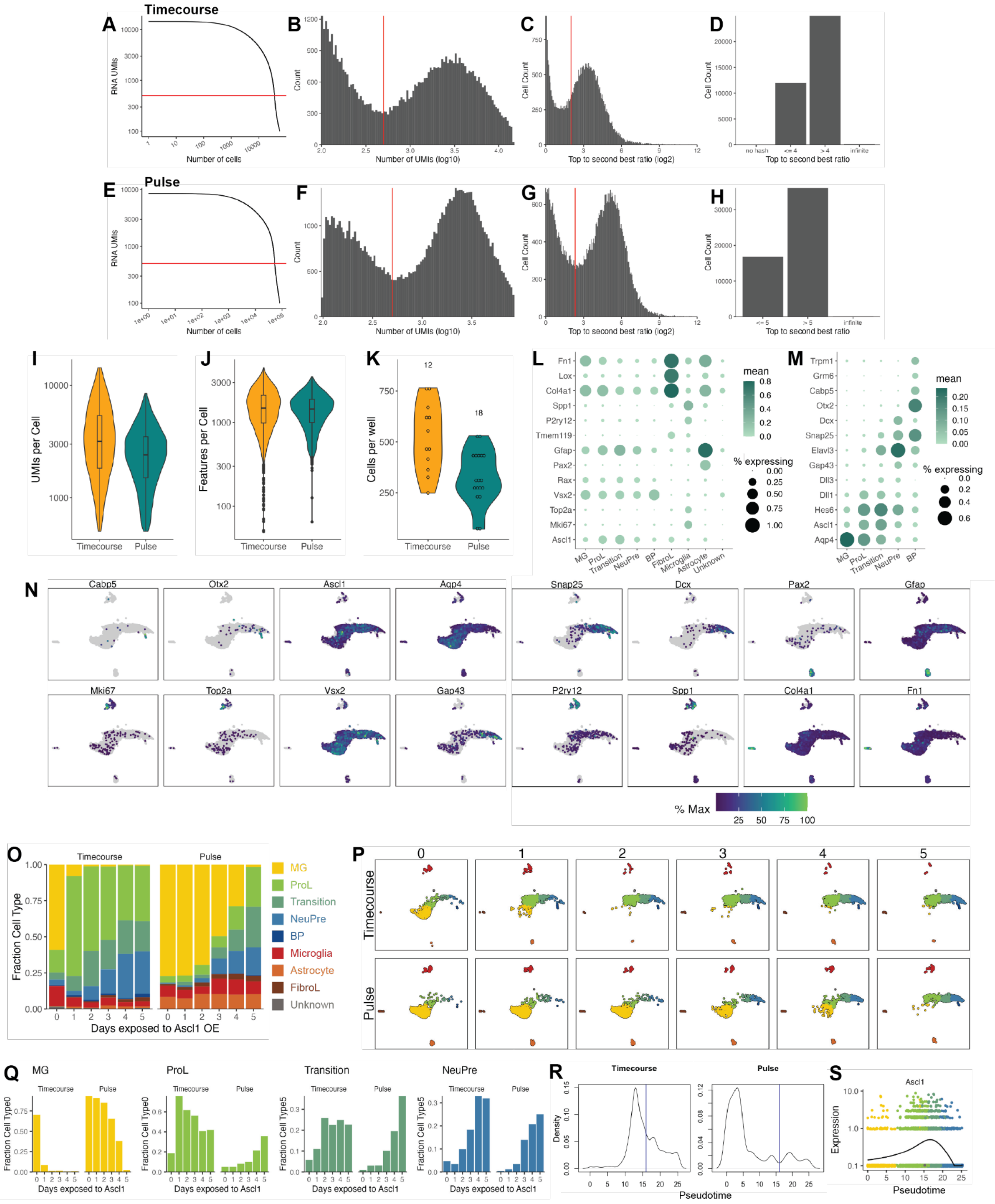

**Fig. S1.** Quality control and cell type annotation for the Timecourse and Pulse experiments. A) Knee plot comparing the number of RNA UMIs recovered to the total number of cells captured for the Timecourse experiment. The red horizontal line is at 500 and indicates the UMI cutoff used in filtering steps. B) Histogram of the number of RNA UMIs (log10) for the Timecourse experiment. The red vertical line is at 500 and indicates the UMI cutoff used in filtering steps. C) Histogram of the hash top_to_second_best_ratio (log2) for the Timecourse experiment. The red vertical line is at 5 and indicates the ratio cutoff used in filtering steps to determine whether a cell was confidently assigned to a condition. D) Bar plot indicating the portion of cells above and below the hash ratio cutoffs for the Timecourse experiment. E) Same as A but for the Pulse experiment. F) Same as B but for the Pulse experiment. G) Same as C but for the Pulse experiment. H) Same as d but for the Pulse experiment. I) Violin plot indicating the number of RNA UMIs recovered per cell for each experiment. J) Violin plot indicating the number of genes/features recovered per cell for each experiment. K) Violin plot quantifying the number of cells recovered per well. The numbers above the plots indicate the number of samples within each experiment. L) Dot plot of the genes used to define the major cell types found in the reprogramming cultures. Dot size indicates the percent of cells that express the gene of interest. The color indicates the log10 mean UMIs per cell. M) Same as L but for the genes used to define cell types derived from MG. N) Gene expression plots of the genes used to annotate cell types. The color for each individual plot is scaled to the maximum expression of that gene. O) Stacked bar plot of the cell type composition across all Ascl1 OE conditions. The colors reflect the cell types. P) UMAP from Figure 1B, but faceted by Ascl1 OE condition. The cells are colored by cell type as in O. Q) The fraction of each cell type at each day of each experiment when considering only the MG, ProL, Transition, and NeuPre cells. R) Cell density plots across pseudotime for the Timecourse and Pulse experiments. The blue vertical line is at 16, the local minimum for pseudotime values > 10 and < 20 in the Pulse experiment. S) Gene expression plots along pseudotime for Ascl1 from the Timecourse experiment only. Each point represents an individual cell’s expression of the indicated gene. The cells are colored by cell type as in O.

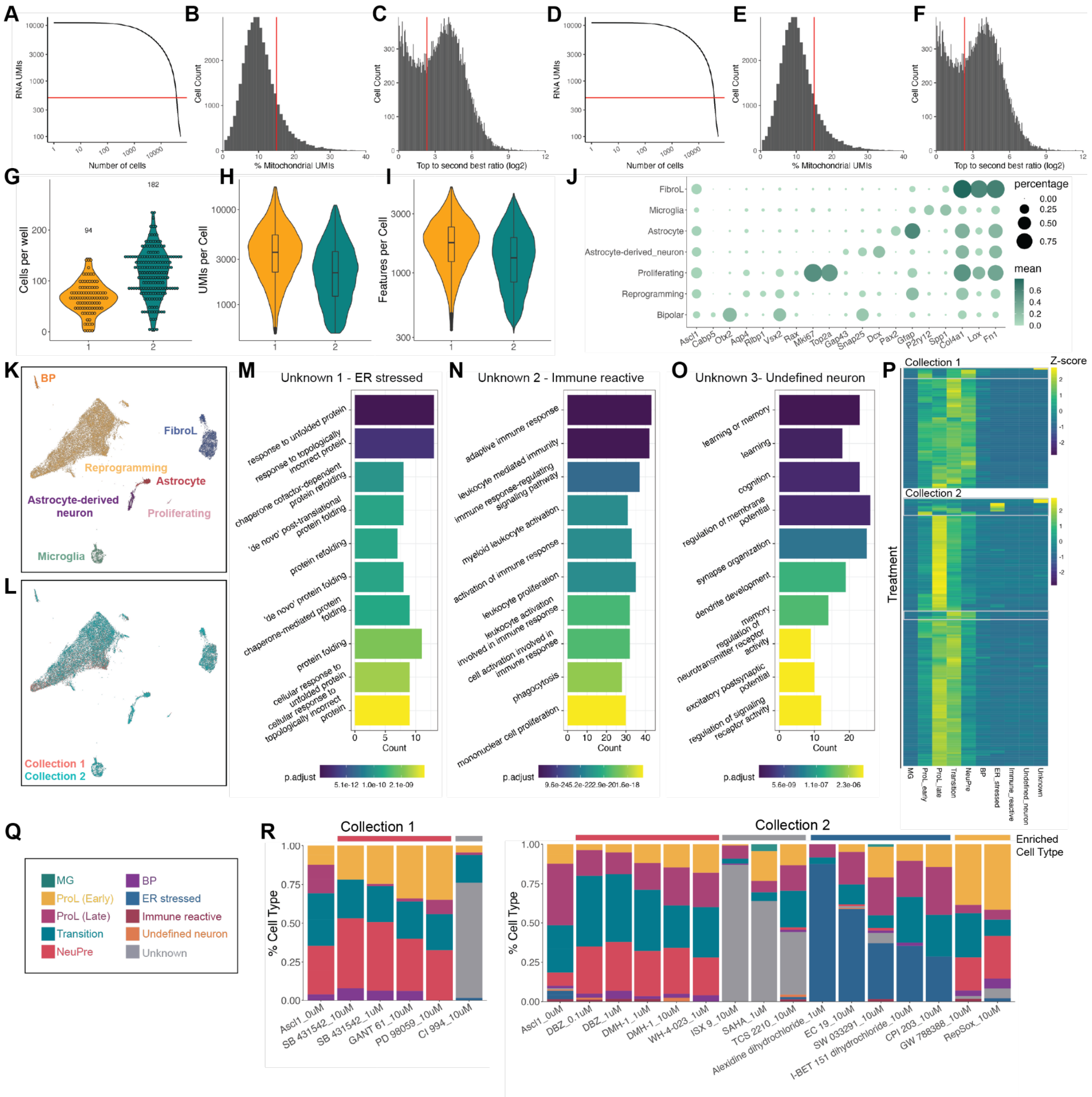

**Fig. S2. Quality control and cell type annotation of small molecule sci-Plex screen. A)** Knee plot comparing the number of RNA UMIs recovered to the total number of cells captured for the Collection 1 experiment. The red horizontal line is at 500 and indicates the UMI cutoff used in filtering steps. **B)** Histogram of the % mitochondrial UMIs for the Collection 1 experiment. The red vertical line at 15% indicates the cutoff used in determining high quality cells. **C)** Histogram of the hash top_to_second_best_ratio (log2) for Collection 1. The red vertical line is at 5 and indicates the ratio above which a cell was determined to be confidently assigned to a single input well. **D)** Same as A but for Collection 2. **E)** Same as B but for Collection 2. **F)** Same as C but for Collection 2. **G)** Violin plot quantifying the number of cells recovered per well. The numbers above the plots indicate the number of samples within each experiment. Each point indicates a treatment well. **H)** Violin plot indicating the number of RNA UMIs recovered per cell for each collection. **I)** Violin plot indicating the number of genes/features recovered per cell for each collection. **J)** Dotplot of the expression patterns of genes used to define cell types from all cells in the screen experiment. Dot size indicates the percent of cells that express the gene of interest. The color indicates the log10 mean UMIs per cell. **K)** Combined UMAP of all the cells captured in Collections 1 and 2. Cells are colored by cell type. **L)** Same UMAP as in K but colored by collection. **M)** GO term enrichment of the genes expressed with high specificity to Unknown 1 in Fig. 2C. The color of the bar reflects the adjusted p-value for each specific GO term. The height of the bar indicates the number of genes in the inputted list of genes that fall into each GO term. **N)** Same as M but for the genes expressed with high specificity to Unknown 2 in Fig. 2C. **O)** Same as M but for the genes expressed with high specificity to Unknown 3 in Fig. 2C. **P)** Heatmap of the cell type abundances across all treatments. Z-scores were calculated by rows. The white boxes indicate treatments that are enriched for Unknown, ER stressed, ProL early, or NeuPre cell types. **Q)** Cell type color legend for R. R) Stacked bar plots representing cell type compositions of each of the treatments highlighted in P.

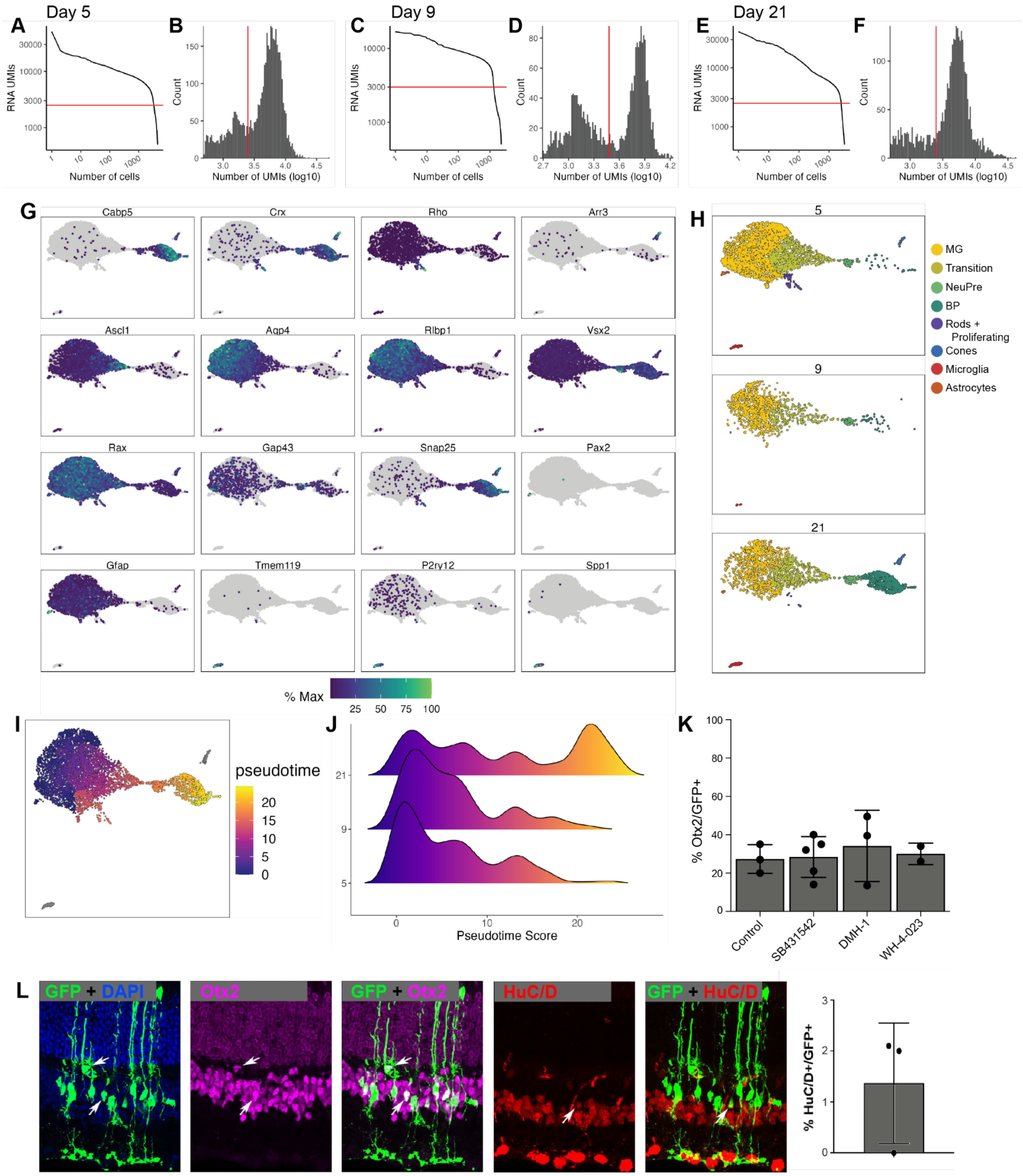


**Fig. S3. Quality control of 10x RNA-seq and immunostaining of *in vivo* reprogrammed neurons. A)** Knee plot comparing the number of RNA UMIs recovered to the total number of cells captured for the Day 5 *in vivo* experiment. The red horizontal line is at 2500 and indicates the UMI cutoff used in filtering steps. **B)** Histogram of the UMIs for the Day 5 *in vivo* experiment. The red vertical line is at 2500 and indicates the UMI cutoff used in filtering steps. **C)** Same as a but for Day 9. The line is at 3000. **D)** Same as B but for Day 9. The line is at 3000. **E)** Same as a but for Day 21. The line is at 2500. **F)** Same as B but for Day 21. The line is at 2500. **G)** UMAP gene expression plots of the genes used to annotate cell types in the reprogramming cells *in vivo* across all timepoints. The color for each individual plot is scaled to the maximum expression of that gene. **H)** UMAP of the cells recovered from the *in vivo* reprogramming faceted by collection day. Cells are colored by cell type. **I)** UMAP of the *in vivo* reprogramming cells colored by pseudotime. **J)** Histograms displaying the frequency of each pseudotime score across *in vivo*

Ascl1 OE durations. **K)** Quantification of the percentage of GFP+ cells that express the bipolar marker Otx2+ in control versus SB431542, DMH-1, or WH-4-023 treatments. None of the treatments showed a significant increase compared to control. **L)** Immunostaining and quantifications of the percentage of GFP+ MG-derived cells that express the HuC/D+ neuronal marker after Metformin treatment.

**Table S1.** Marker genes used for cell type annotation.

| EXPERIMENT | CELL TYPE | MARKERS | | | | | | |
|---|---|---|---|---|---|---|---|---|
| TC/PULSE | MG | Aqp4 | | | | | | |
| TC/PULSE | ProL | Ascl1 | ↓Aqp4 | Hes6 | Dll1 | Dll3 | | |
| TC/PULSE | Transition | high Ascl1 | ↓Aqp4 | ↑Snap25 | ↑Dcx | Hes6 | Dll1 | Dll3 |
| TC/PULSE | NeuPre | Snap25 | Dcx | Gap43 | Hes6 | Dll3 | Elavl3 | |
| TC/PULSE | BP | Otx2 | Cabp5 | ↑Snap25 | ↓Hes6 | ↓Dll3 | Trpm1 | Grm6 |
| TC/PULSE | Astrocyte | Gfap | Pax2 | | | | | |
| TC/PULSE | Microglia | Spp1 | P2ry12 | | | | | |
| TC/PULSE | FibroL | Fn1 | Lox | Col4a1 | | | | |
| TC/PULSE | Unknown | | | | | | | |
| SCREEN_ALL | Reprogramming | Vsx2 | Aqp4 | Rlbp1 | | | | |
| SCREEN_ALL | Bipolar | Otx2 | Cabp5 | ↑Snap25 | ↓Hes6 | ↓Dll3 | Trpm1 | Grm6 |
| SCREEN_ALL | Astrocytes | Gfap | Pax2 | | | | | |
| SCREEN_ALL | Microglia | Spp1 | P2ry12 | | | | | |
| SCREEN_ALL | FibroL | Fn1 | Lox | Col4a1 | | | | |
| SCREEN_ALL | Astrocyte-derived neuron | Gap43 | Dcx | Snap25 | ↓Vsx2 | | | |
| SCREEN_ALL | Proliferating | Mki67 | Top2a | Col4a1 | Fn1 | Lox | Vsx2 | Aqp4 |
| SCREEN_MG | MG | Aqp4 | Apoe | | | | | |
| SCREEN_MG | ProL (early) | Apq4 | Gfap | Apoe | Hes5 | Fosb | Ascl1 | |
| SCREEN_MG | ProL (late) | Ascl1 | Fn1 | Hes5 | | | | |
| SCREEN_MG | Transition | Ascl1 | Gfap | Snap25 | | | | |
| SCREEN_MG | NeuPre | Gap43 | Snap25 | Dcx | | | | |
| SCREEN_MG | BP | Snap25 | Vsx2 | Cabp5 | Otx2 | Grik1 | | |
| SCREEN_MG | ER stressed | Hspb1 | Trib3 | | | | | |
| SCREEN_MG | Immune reactive | Pik3ap1 | Mpeg1 | | | | | |
| SCREEN_MG | Undefined neuron | Otx1 | Neto1 | | | | | |
| SCREEN_MG | Unknown | Slc5a5 | Perp | | | | | |
| IN_VIVO | MG | Aqp4 | Rlbp1 | | | | | |
| IN_VIVO | Transition | Aqp4 | Rlbp1 | Ascl1 | | | | |
| IN_VIVO | NeuPre | Otx2 | Nrxn3 | | | | | |
| IN_VIVO | Bipolar | Snap25 | Cabp5 | Syt1 | Nrxn3 | | | |
| IN_VIVO | Cones | Arr3 | Syt1 | Snap25 | | | | |
| IN_VIVO | Rods and Proliferating Cells | Rho | Nrl | Mki67 | Top2a | | | |
| IN_VIVO | Astrocyte | Pax2 | Gfap | | | | | |
| IN_VIVO | Microglia | Tmem119 | P2ry12 | Spp1 | | | | |

**Dataset S1 (separate file).** Genes differentially expressed across pseudotime.

**Dataset S2 (separate file).** Fold change calculations for small molecules used from Tocriscreen Stem Cell Library.

**Dataset S3 (separate file).** Genes used and GO annotations in unknown clusters.
